# Reproduction, seasonal morphology, and juvenile growth in three Malagasy fruit bats

**DOI:** 10.1101/2021.10.28.466299

**Authors:** Angelo Andrianiaina, Santino Andry, Anecia Gentles, Sarah Guth, Jean-Michel Héraud, Hafaliana Christian Ranaivoson, Ny Anjara Fifi Ravelomanantsoa, Timothy Treuer, Cara E. Brook

## Abstract

The island nation of Madagascar is home to three endemic species of Old World Fruit Bat in the family Pteropodidae: *Pteropus rufus, Eidolon dupreanum*, and *Rousettus madagascariensis*, all three of which are IUCN Red Listed under some category of threat. To inform conservation efforts to model population viability for these threatened species, as well understand the mechanisms underpinning persistence of several potentially zoonotic pathogens hosted by these bats, we here define the seasonal limits of a staggered annual birth pulse across the three species. Our field studies in central-eastern Madagascar indicate that this annual birth pulse takes place in September/October for *P. rufus*, November for *E. dupreanum*, and December for *R. madagascariensis*. Juvenile development periods vary across the three Malagasy pteropodids, resulting in near-synchronous weaning of pups for all species in late January-February at the height of the fruiting season for Madagascar, a pattern characteristic of most mammalian frugivores on the island. We here document the size range in morphological traits for the three Malagasy fruit bat species; these traits span the range of those known for pteropodids more broadly, with *P. rufus* and *E. dupreanum* among the larger of recorded species and *R. madagascariensis* among the smaller. All three species demonstrate subtle sexual dimorphism in observed traits with larger-bodied males vs. females. We explore seasonal variation in adult body condition by comparing observed body mass with body mass predicted by forearm length, demonstrating that pregnant females add weight during staggered gestation periods and males lose weight during the nutritionally-deficit Malagasy winter. Finally, we quantify forearm, tibia, and ear length growth rates in juvenile bats, demonstrating both faster growth and more protracted development times for the largest *P. rufus* species. The longer development period for the already-threatened *P. rufus* further jeopardizes this species’ conservation status as human hunting of bats for subsistence is particularly detrimental to population viability during reproductive periods. The more extreme seasonal variation in the mass to forearm relationship for *P. rufus* may also modulate immune function, an important consideration given these bats’ roles as reservoir hosts for several high profile viral families known to cause severe disease in humans. Our work highlights the importance of longitudinal field studies in collecting critical data for mammalian conservation efforts and human public health alike.

---

The Old World Fruit Bat family Pteropodidae, known colloquially as the ‘flying foxes,’ makes up one of the most endangered groups of mammals on Earth, with some 35% of species either extinct or threatened with extinction, a proportion almost three times higher than that reported (12%) for all other bat families combined (Species IUCN Red List Threat. 2018). Fruit bats experience disproportionate rates of persecution, likely as a result of their propensity for small island endemism (Jones et al. 2009) and their large sizes (fruit bat wingspans can reach up to two meters in the case of *Pteropus vampyrus*, the world’s largest bat; Corbet and Hill 1992), which make them targets for the bushmeat trade (Craig et al. 1994; Brooke 2002; Oleksy et al. 2003; Jenkins and Racey 2008; Kamins et al. 2011; Openshaw et al. 2016; Peel et al. 2017). Pteropodid bats offer critical services to surrounding ecosystems, playing important roles in the pollination and seed dispersal of numerous plant species across the Old World, particularly in island ecosystems often depauperate in other frugivores (McConkey and Drake 2006; Kunz et al. 2011).

Madagascar is one such island ecosystem recognized for its unusually depauperate frugivorous fauna (Dewar and Richard 2007). Primates (lemurs), rather than birds, are considered the primary seed dispersers on the island (Langrand 1990; Wright et al. 2011), in contrast to otherwise comparable tropical ecosystems in the New World (Terborgh 1983, 1986). In addition to lemurs, Madagascar is home to three endemic species of frugivorous bats from the family Pteropodidae—*Pteropus rufus, Eidolon dupreanum*, and *Rousettus madagascariensis—*all of which are known to pollinate flowers and disperse seeds from both native Malagasy and exotic plants (Bollen and Elsacker 2002; Andriafidison et al. 2006; Long and Racey 2007; Picot et al. 2007; Andrianaivoarivelo et al. 2011; Oleksy et al. 2015, 2017). Importantly, *E. dupreanum* may be the only extant pollinator of the endangered, endemic Malagasy baobab, *Adansonia suarezensis* (Andriafidison et al. 2006).

Despite their ecosystem value, Madagascar’s fruit bats are heavily persecuted. All three species are consumed across the island as a source of human food (Oleksy et al. 2003; Jenkins and Racey 2008; Cardiff et al. 2009; Randrianandrianina et al. 2010; Golden et al. 2014; Fernández-Llamazares et al. 2018; Brook et al. 2019a), and *P. rufus*, the largest and most heavily hunted, is sometimes targeted in response to its largely inaccurate characterization as a predator of human fruit crops (Raharimihaja et al. 2016). Respectively, *P. rufus, E. dupreanum*, and *R. madagascariensis* are currently IUCN Red-listed as ‘Vulnerable,’ ‘Vulnerable,’ and ‘Near-Threatened’ species (Species IUCN Red List Threat. 2018), though recent population viability analyses suggest that *P. rufus*, in particular, may be experiencing more severe population declines than have been previously reported (Brook et al. 2019a). Bats are reservoir hosts for a majority of the world’s most virulent zoonotic viruses (Guth et al. 2019, 2021), as well as hosts for coronaviruses ancestral to the recently emerged SARS-CoV-2 (Zhou et al. 2020; Temmam et al. 2021). Globally, anti-bat sentiments have been on the rise as a result of the COVID-19 pandemic (Rocha et al. 2020); though no specific instances of COVID-related persecution have yet been documented for the Malagasy fruit bats, all three species are known to host potentially zoonotic pathogens (Iehlé et al. 2007; Razafindratsimandresy et al. 2009; Reynes et al. 2011a; Wilkinson et al. 2012a; Brook et al. 2015, 2019b; Razanajatovo et al. 2015; Ranaivoson et al. 2019), posing risks that negative public reactions may arise in the future.

Previous work suggests that roost population sizes and survival rates vary across the year for these three species (Brook et al. 2019a; Noroalintseheno Lalarivoniaina et al. 2019). Temporal fluctuations in nutritional status may alter bat immune responses, thus influencing pathogen dynamics (Brook et al. 2019b), as well as modulate bats’ vulnerability to seasonally variable hunting pressures (Brook et al. 2019a). All three Malagasy fruit bats are thought to reproduce seasonally in species-specific annual birth pulses (MacKinnon et al. 2003; Brook et al. 2019a). Documentation of the timing of these birth pulses for Malagasy fruit bats is important for understanding their vulnerability to seasonally-varying population pressures: previous work describes how seasonal variation in hunting pressure for Malagasy lemurs poses elevated risks to species when directly overlapping their annual birth pulse (Brook et al. 2018).

In addition to its importance for conservation efforts to quantify fruit bat population viability, defining the temporal limits of each fruit bat species-specific birth pulse is essential to understanding the mechanisms which underpin the maintenance and persistence of numerous infectious agents that these bats host (Iehlé et al. 2007; Razafindratsimandresy et al. 2009; Reynes et al. 2011b; Wilkinson et al. 2012b; Brook et al. 2015, 2019b; Ranaivoson et al. 2019). Isolated *E. helvum* populations on islands off the west coast of Africa have been shown to support circulation of potentially zoonotic henipaviruses at population sizes well below the established critical community size for closely-related paramyxoviruses in other systems (Bartlett 1957, 1960; Swinton et al. 1998; Peel et al. 2012). Some work has suggested that seasonally-staggered births allowing for a protracted introduction of juvenile susceptibles into the host population could play a role in pathogen persistence in these systems (Peel et al. 2013, 2014; Hayman 2015).

We sought to expand existing knowledge of seasonal variation in the reproductive calendar and nutritional status of all three Malagasy fruit bat species, to facilitate future conservation assessments and studies aimed at deciphering the dynamics of bat-hosted infections. In particular, we aimed to (a) quantify life history traits needed for population modeling for these species, (b) document seasonal variation in morphometrics and body conditions for these bats, and (c) calculate juvenile growth rates throughout the post-reproductive period. Our work emphasizes the importance of longitudinal field studies in accurately describing the ecology of frugivorous bats.

## Materials and Methods

### Study periods and sites

Field studies were carried out between 2013 and 2020 in part with previously published work examining population viability and the dynamics of potentially zoonotic infections in Malagasy fruit bats (Brook et al. 2015, 2019b; a; Ranaivoson et al. 2019). Bats were captured periodically throughout each year, with sampling spanning all months and all seasons (dry, wet, shoulder), interspersed with some gaps in temporal continuity. Captures took place in several regions of Madagascar: (1) Ankarana National Park in the northwest (−12.9S, 49.1E), (2) Makira Natural Park in the northeast (−15.1S, 49.6E), (3) Mahabo forest in the center-west (−20.5S, 44.7E), and (4) several sub-localities of the Moramanga District in the center-east, including: the fragmented forests of Ambakoana (−18.5, 48.2), Mangarivotra (−18.3S, 48.2E), Marotsipohy (−18.4S,48.1E), Marovitsika (−18.8S,48.1E), Lakato (−19.2S, 48.4E), and Mahialambo (−18.1S, 48.2E), the special reserves of Angavokely (−18.9S, 47.8E) and Angavobe (−18.9S), 47.9E, and the new protected area of Maromizaha (−18.9S, 48.5E).

### Netting

Mist nets were deployed from 6:00 p.m. to midnight and from 3:00 a.m. to 8:00 a.m. around roosting or feeding sites of *P. rufus, E. dupreanum* and *R. madagascariensis* and monitored continuously. Captured bats were placed in individual clean cloth bags while awaiting processing for infectious disease studies, as has been previously described (Brook et al. 2015, 2019b; Ranaivoson et al. 2019). For each sampling session, we conducted between 1 and 10 nights of netting, ending sessions early when 30 individuals of each species present at the site were captured. Upon capture, all bats were weighed (in grams) with a Pesola scale attached to the cloth bag and forearm, tibia, and ear were measured with a caliper or tape measure (in mm). Bats were classed by sex and age (juvenile vs. adult) and, for females, reproductive class (non-reproductive, pregnant, lactating). For females captured approximately within the period of possible gestation for each species, abdominal palpitation was used to determine whether or not females were pregnant. All raw data used in this study are accessible in our open-access GitHub repository at: github.com/brooklabteam/Mada-Bat-Morphology.

This study was carried out in strict accordance with research permits obtained from the Madagascar Ministry of Forest and the Environment (permit numbers 251/13, 166/14, 075/15, 258/16, 170/18, 019/18, 170/18, 007/19, 14/20) and under guidelines posted by the American Veterinary Medical Association. All field protocols employed were pre-approved by the Princeton University and UC Berkeley Institutes for Animal Care and Use Committees (respectively, IACUC Protocol #1926 and ACUC Protocol # AUP-2017-10-10393), and every effort was made to minimize discomfort to animals.

### Literature review

To place our Malagasy bats in a broader context, we compiled information from the literature concerning the morphology of other bats in the family Pteropodidae. From the ‘Bat Species of the World’ database (Simmons and Cirranello 2020), we compiled a list of 201 previously described pteropodid species, then searched GoogleScholar and Web of Science for any records documenting the mass, forearm, tibia, and ear length of each species. We only collected records that were sex-specific, and where possible, we documented the sample size from which those records were derived, if reported as an average. In cases where no sample size was reported, we assumed sample size to be one individual. All raw data and references are accessible in our open-access GitHub repository at: github.com/brooklabteam/Mada-Bat-Morphology.

### Statistical analysis

Data analysis was performed using R v.4.0.3 (R Core Team, 2020). All raw data and corresponding code for these analyses can be accessed in our GitHub repository. Additional details of statistical output are compiled in supplementary tables in Appendix 2. First, we aimed to define the seasonal limits of the reproductive calendar for each of the three Malagasy fruit bat species. To this end, we restricted our analyses to our most complete cross-species time series from roost sites in the District of Moramanga, Madagascar and queried the data subset for the following metrics, unique for each species: (a) the earliest calendar day on which a pregnant female was observed, (b) the earliest calendar day on which a juvenile was observed, and (c) the latest calendar day on which a lactating female was observed. Metrics (a) and (b) corresponded to the date limits of gestation for each species, while metrics (b) and (c) corresponded to the date limits of lactation for each species. Because fruit bats of many species are known to delay embryonic implantation and fetal development for months after fertilization (Mutere 1967; Heideman 1988; Heideman and Powell 1998; Meenakumari and Krishna 2005), we assumed that abdominal palpitation to determine reproductive status in the field would likely miss early-stage pregnancies in the three Malagasy species. To this end, we additionally searched the literature for records of gestation length in closely-related pteropodids to compare against our records of observed gestation in Malagasy species.

We next sought to document morphological variation in adult *P. rufus, E. dupreanum*, and *R. madagascariensis*, as compared with other bats in family Pteropodidae. To this end, we calculated the sex-specific median and interquartile range of reported measurements of mean tibia and ear length (in mm) for adult pteropodids globally, as well as the range of values recorded for individuals within our dataset. To investigate any potential sexual dimorphism in our dataset, we compared mean forearm, tibia, and ear length for male vs. female distributions across the global dataset and the three Malagasy species using *Welch’s 2-sample t-tests* for independent distributions of unequal sample size.

We additionally compared the relationship between sex-specific forearm length and mass for adult pteropodids surveyed in the literature against the ranges recorded in our own field data for the three Malagasy species. We first fit a linear regression to log_10_-transformed values for both forearm length (predictor variable) and mass (response variable), separated by sex, both to species-level averages for pteropodids globally and to individual datapoints for adults of the three Malagasy species.

Next, we explored seasonal variation in the relationship between adult body mass and forearm length within our Malagasy fruit bat field data. To facilitate this analysis, we refit a composite linear regression using log_10_-transformed values for forearm length as predictors of log_10_-transformed values for mass. We included a fixed effect of bat species as an additional predictor to control for differences in skeletal structure:mass ratios across the different species, then calculated the residual of each individual’s observed mass in the data against that predicted from the regression. This generated a body condition index metric for bats: individuals with positive mass:forearm residuals corresponded to those with higher masses than predicted by body size (broadly indicative of better nutritional condition), while individuals with negative mass:forearm residuals corresponded to those with lower masses than predicted by body size (broadly indicative of poorer nutritional condition). Because all individuals included in these analyses were adults, and previous work indicates that up to 96% of reproductively mature fruit bats give birth annually (Hayman et al. 2012; Brook et al. 2019a), we assumed all females observed during the defined gestation period for each species to be pregnant, regardless of reproductive class recorded from abdominal palpitation.

To assess seasonal variation in body condition, we restricted our analysis to data collected from the longitudinally-monitored Moramanga sites only, and fit a generalized additive model (GAM), using the mgcv package in R (Wood 2001), to the seasonal time series of mass:forearm residual, separately across each discrete species-sex subset of the data. We modeled mass:forearm residual as the response variable predicted by day of year as a cyclic cubic (“cc”) spline, with the number of smoothing knots (“k”) fixed at seven, as recommended by the package author (Wood 2001). Cyclic cubic splines can be used to capture annual seasonality, as the seasonal smoother on January 1 is modeled as a continuation from December 31. Because some previous work has questioned the effectiveness with which body condition indices represent bat nutritional status (McGuire et al. 2018), we computed an additional set of supplementary GAMs which included the predictor variables of day of year (as a cyclic cubic smoothing spline) and forearm length (as a random effect) against the response variable of mass (in grams) for both sexes and all three species.

Finally, we explored juvenile growth rates for forearm, tibia and ear across all three Malagasy fruit bat species, calculating the age in days since birth of each juvenile bat in our dataset with “day 0” set equal to the first date of an observed juvenile in the dataset for each species, as described above, up to one year of life (day 365). Using GAMs, we then modeled the response variables of forearm length, tibia length, and ear length against the smoothing predictor of age in days, using a thinplate smoothing spline (“tp”) with the number of smoothing knots fixed again at seven. After fitting each model, we then calculated the age-varying derivative of each fitted curve using the ‘gratia’ package in R to facilitate comparison of growth rates across different species and morphological features.

## Results

### Field captures

In total, 2160 fruit bats were captured and processed between August 2013 and March 2020 (**Fig. 1A**). The majority of bats (n=1700) were captured in roost sites located in the District of Moramanga in central-eastern Madagascar (*P. rufus* n=316; *E. dupreanum* n=732; *R. madagascariensis* n=652), followed by Ankarana National Park in the northwest (n=380; *E. dupreanum* n= 172; *R. madagascariensis* n =208), Makira Natural Park in the northeast (n=47; *P. rufus* n=15; *R. madagascariensis* n=32), and Mahabo forest in the center-west (n= 33; *P. rufus* n=19; *E. dupreanum* n=14) (**Table S1**).

**Fig. 1.**
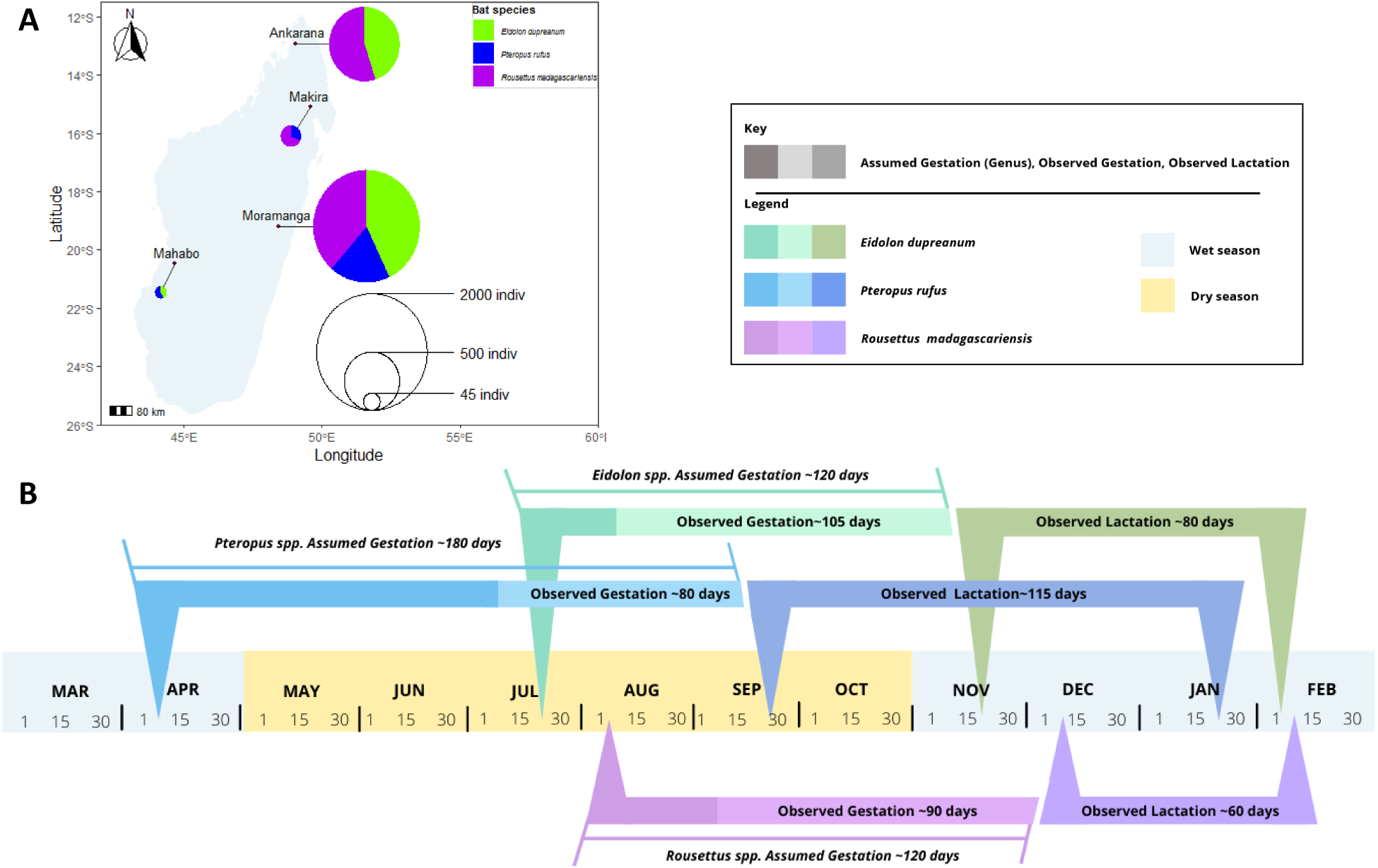
B. Map of field sites and distribution of bat captures for *P. rufus, E. dupreanum*, and *R. madagascariensis* in Madagascar. Pie size corresponds to total bats captured at each site: 1700 in the District of Moramanga (*P. rufus* n=317; *E. dupreanum* n=732; *R. madagascariensis* n=653), 380 in Ankarana National Park (*E. dupreanum* n= 172; *R. madagascariensis* n =208), 47 in Makira Natural Park (*P. rufus* n=15; *R. madagascariensis* n=32), and 33 in Mahabo forest (*P. rufus* n=19; *E. dupreanum* n=14). **B**. Gestation and lactation periods across the three Madagascar fruit bat species, calculated from the field data (observed) and reported in the literature (assumed). Respectively, for *P. rufus, E. dupreanum*, and *R. madagascariensis*, observed gestation begins on: July 7, August 3, and September 11; birth occurs on: September 29, November 16, and December 12; and lactation ceases on: January 21, February 2, and February 19 (Table S1).

### Fruit bat reproductive calendars

Longitudinal data collected in the District of Moramanga allowed us define the seasonal limits of a single annual reproduction event for all three fruit bat species (**Fig. 1B**). We calculated the earliest calendar day on which a pregnant female was observed, respectively, for *P. rufus, E. dupreanum*, and *R. madagascariensis*, as July 7, August 3, and September 11; the earliest calendar day on which a juvenile was observed as September 29, November 16, and December 12; and the latest calendar day on which a lactating female was observed as January 21, February 2, and February 19 (**Table S1**). These dates allowed us to define the approximate duration of the observed gestation and lactation period for each species (observed gestation: *P. rufus* = ∼80 days, *E. dupreanum=* ∼105 days, and *R. madagascariensis =* ∼90 days; observed lactation: *P. rufus* = ∼115 days, *E. dupreanum=* ∼80 days, and *R. madagascariensis =* ∼60 days). Because gestation was observed through abdominal palpitation in the field, we presumed that early stage pregnancies for all three species might not be visible. To account for this, we compared our observed gestation period for all three fruit bat species against that which has been previously described for closely-related species: *Pteropus alecto, Pteropus policephalus*, and *Pteropus scapulatus* (sister species to *P. rufus*) demonstrate a ∼180 day gestation period on the Australian continent (McIlwee and Martin 2002), while *Eidolon helvum* (sister species to *E. dupreanum*) and *Rousettus aegyptiacus* (sister species to *R. madagascariensis*) both demonstrate gestation periods of ∼120 days on the African continent (Odukoya et al. 2008; Barclay and Jacobs 2011). Extension of the gestation period for the three Malagasy species back in time from the birth pulse to match those recorded for sister species elsewhere would place the mating period for *P. rufus* in the month of April, for *E. dupreanum* in the month of July, and for *R. madagascariensis* in the month of August. These estimates of mating period are consistent with previous reporting for *P. rufus* (Long and Racey 2007) and *R. madagascariensis* (Noroalintseheno Lalarivoniaina et al. 2019); to our knowledge, no previous records of the reproductive calendar for *E. dupreanum* have been published.

In sum, we observed the longest gestation and lactation period for *P. rufus*, which births first of the three Malagasy fruit bat species, followed by *E. dupreanum*, and *R. madagascariensis*, in order of decreasing body size. Despite differences in the timing and duration of gestation, lactating mothers for all three species weaned pups around the same time of the year (∼late January – February), at the onset of peak fruit abundance in the hot-wet season in the District of Moramanga.

### Morphological patterns

After searching the literature, we successfully compiled adult mass records from 103 pteropodid species for females and 106 species for males; adult forearm records from 146 species for females and 140 species for males; adult tibia records from 64 species for females and 64 species for males; and adult ear length records from 101 species for females and 99 species for males. We compared these records against morphological patterns witnessed in our own longitudinally-collected field data.

For Malagasy fruit bats, we observed large differences in adult morphology across species and more subtle differences by sex. The Malagasy fruit bat species ranked in size, from largest to smallest: *P. rufus, E. dupreanum*, and *R. madagascariensis*, with the size ranges of each species roughly spanning the range in ear length, tibia length, and forearm length captured across mean values for all non-Malagasy pteropdid bats surveyed in the literature (**Fig. 2A-C**). Specific morphological ranges for tibia length and forearm length matched the size distributions of the three Malagasy species, scaling downward from *P. rufus* to *E. dupreanum* to *R. madagascariensis*. For ear lengths, *P. rufus* and *E. dupreanum* distributions were largely overlapping, while *R. madagascariensis* were smaller; species-specific interquartile ranges for each morphological trait are summarized in **Table S2**. Global data for comparison roughly approximated the range spanned from the *R. madagascariensis* minimum to the *P. rufus* maximum, with the median falling in between that of *R. madagascariensis* and *E. dupreanum* across all three metrics.

**Fig. 2.**
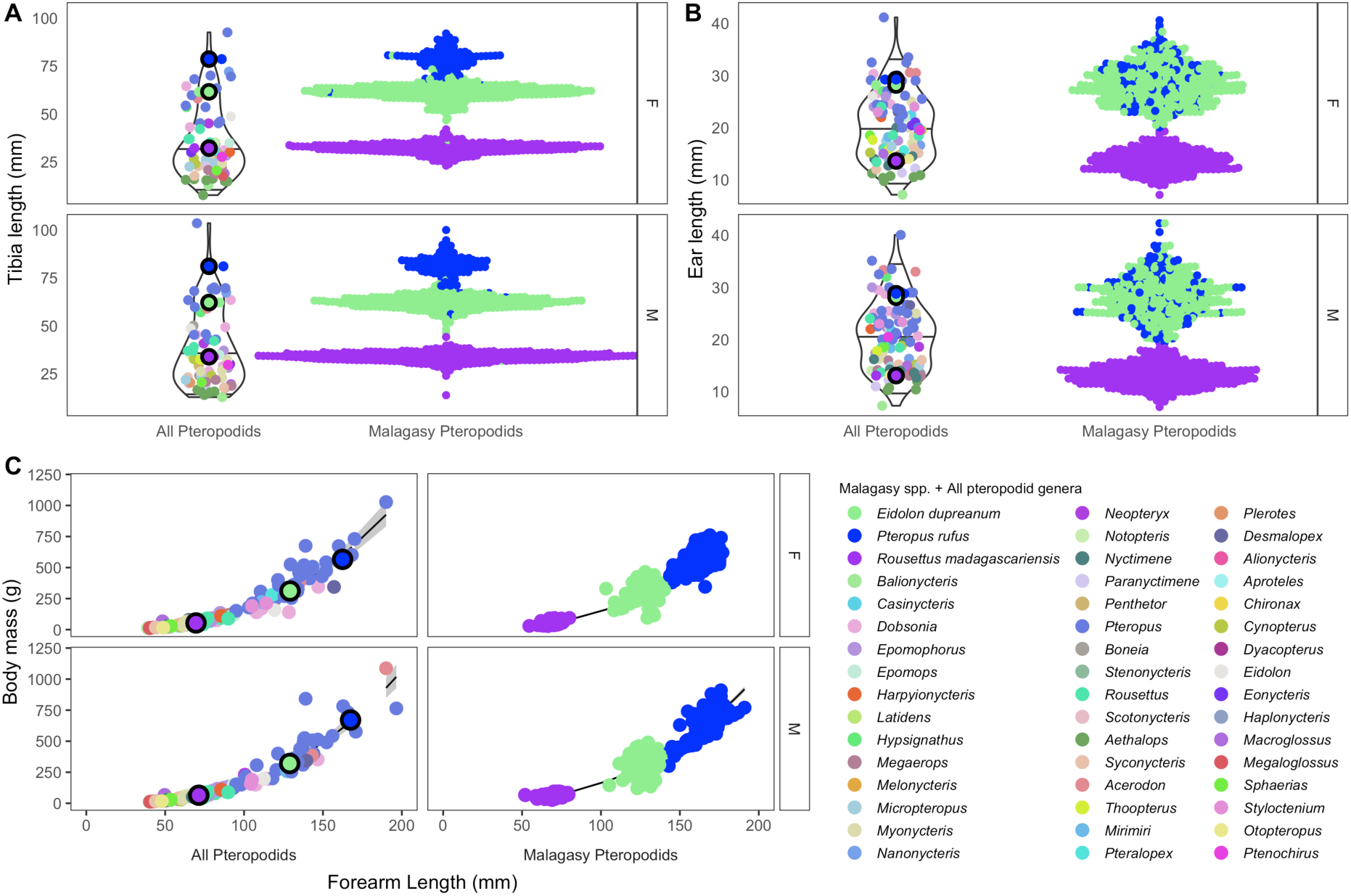
**A**. Tibia, and **B**., ear length across fruit bat species from the literature (left) and from our Madagascar field data (right), colored by genera according to legend; data are grouped by sex (upper=females, lower=males). Violin plots show range and 25, 50, and 75% quantiles for each distribution. **B**. Linear regression of log_10_ body mass (in grams, y-axis) by log_10_ forearm length (in mm, x-axis) across pteropodids from the literature (left) and from our Madagascar field data (right), colored by genera according to legend; data are grouped by sex (upper=females, lower=males). Solid line corresponds to predictions from the fitted model (R^2^: All Pteropodids, M= .96, F=.95; Malagasy Pteropodids, M=.96, F=.97). Data are summarized in Table S2, S3.

*Welch’s 2-sample t-test* comparisons indicated that length distributions for tibia and forearm length were significantly longer in adult males vs. females for both *P. rufus* and *R. madagascariensis* (*p<0*.*001;* Table S2; **Fig. S1**). For *E. dupreanum*, only tibia length was different between the sexes, with males again larger than females (*p<0*.*029)*. Ear lengths showed sexual dimorphism only in *R. madagascariensis* bats, for which observed female ear lengths were actually larger than those of males. Nonetheless, given that both tibia and forearm length were larger in *R. mdagascariensis* males vs. females, we conclude that all three Malagasy fruit bat species demonstrated slight sexual dimorphism characterized by larger-bodied males and smaller females.

Linear regressions of log10 body mass as predicted by log10 forearm length for both all-pteropodid and Malagasy-specific datasets, separated by sex, demonstrated a good fit to the data with *R*^*2*^ values > .95. Roughly comparable slopes across all four models indicated 20-30 fold increases in bat mass (in grams) corresponding to every 10-fold increase in forearm length (in mm) across all species and sexes (Fig 2C; **Table S3**).

### Seasonality of mass:forearm relationships

Restricting our analyses to the Madagascar field data only, we refit the regression of mass:forearm length across both sexes, incorporating bat species as a second fixed predictor of mass (**Table S4; Fig. S2**), then computed mass:forearm residuals from the resulting regression models for each individual. We explored seasonal variation in these residuals from the longitudinally-monitored Moramanga site only, using GAMs. GAM results indicated significant seasonality in bat body condition for both male and female subsets of the *P. rufus* and *E. dupreanum* data and for the female subset of the *R. madagascariensis* data (*p-value<0*.*001;* **Table S5**). Only male *R. madagascariensis* demonstrated no seasonal variation in mass:forearm residual. Finally, we plotted the GAM-predicted mass for each species and sex across, respectively, the reproductive and nutritional calendars for female and male fruit bats of the three Malagasy species (**Fig. 3**). As expected, we observed a seasonal peak in adult female mass:forearm which overlapped the staggered period of observed gestation for each species from Fig. 1, followed by a deficit overlapping the corresponding, species-specific lactation period. These results supported our assumption that the vast majority of female bats in our dataset could be considered reproductive.

**Fig. 3.**
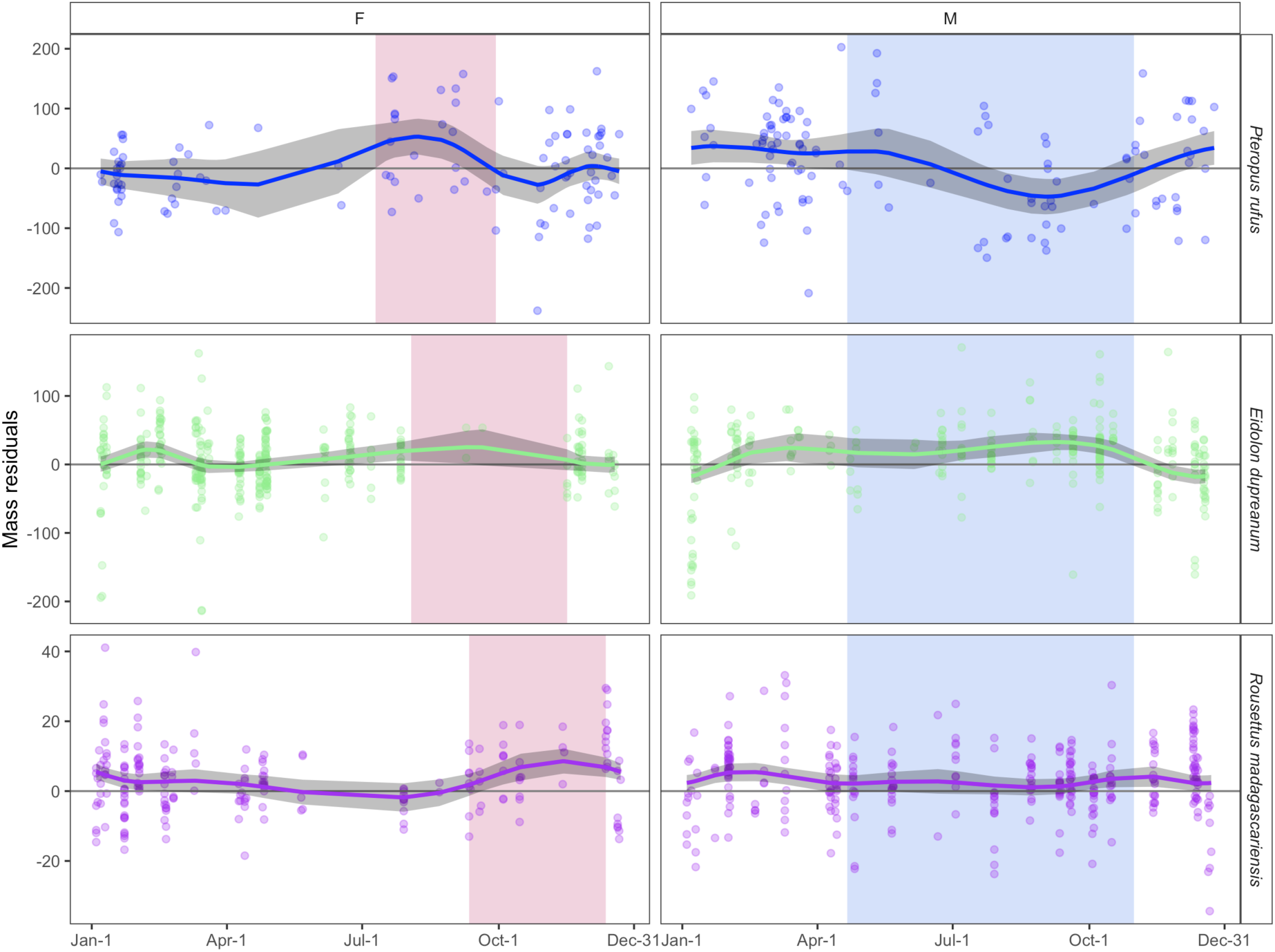
Seasonal variation in mass:forearm residual by sex (females = left, males = right) and species. Raw data are shown as open circles with prediction from fitted GAM model as solid line; 95% confidence intervals by standard error are shown by shading in gray (Table S5). For female plots, pink shading corresponds to the species-specific gestation period; for male plots, blue shading corresponds to the winter dry season in Madagascar.

We also observed a less extreme mass deficit which overlapped the resource-poor winter for male *P. rufus* and *E. dupreanum* but occurred earlier in the season for *E. dupreanum* than for *P. rufus*. Notably, GAMs for *R. madagascariensis* males, which showed no significant seasonality in body condition, predicted positive mass:forearm residuals across the entire calendar year, suggesting that bats in the Moramanga site had high mass:forearm ratios, as compared with those across the entire dataset (sites outside of Moramanga were not included in seasonal analyses). These results are logical, considering that, outside of Moramanga, *R. madagascariensis* were predominantly captured in Ankarana National Park, an arid, desert environment where bats are much more likely to experience food stress. Finally, supplementary GAMs used to model seasonal mass directly demonstrated comparable results to patterns for mass:forearm residuals across all three species and both sexes, with female masses peaking across gestation and male masses declining through the winter season (**Fig. S3**).

### Juvenile growth rates

In our final analysis, we compared juvenile growth rates in forearm, tibia, and ear length across all three Malagasy fruit bat species in the first year of life. GAMs fitted to the response variable of each morphological trait demonstrated highly significant smoothing predictors of days since birth across all three metrics and all three species (**Fig. 4; Table S6**). Quantification of the derivative of each fitted GAM across the range of observed days since birth allowed us to compare growth rates across traits and species: in general, we observed the largest slopes, corresponding to the fastest growth rates for forearm lengths, then tibia lengths, and finally, ear lengths of all three species. *P. rufus* grew at the fastest rate (largest slope in growth curve) for all three morphological traits, followed by *E. dupreanum* and then *R. madagascariensis*. Despite faster growth rates, as the largest of the three species, *P. rufus* also demonstrated the most protracted development phase, approaching adult size (10-day average slope for forearm growth <.1) approximately six months after birth (180 days), as compared to two months (53 days) for *E. dupreanum* and 2.5 months (81 days) for *R. madagascariensis*. Species- and metric-specific growth rates from our fitted GAMs across the first year of life are summarized in **Table S7**; all raw values are available on our open-access GitHub repository.

**Fig. 4.**
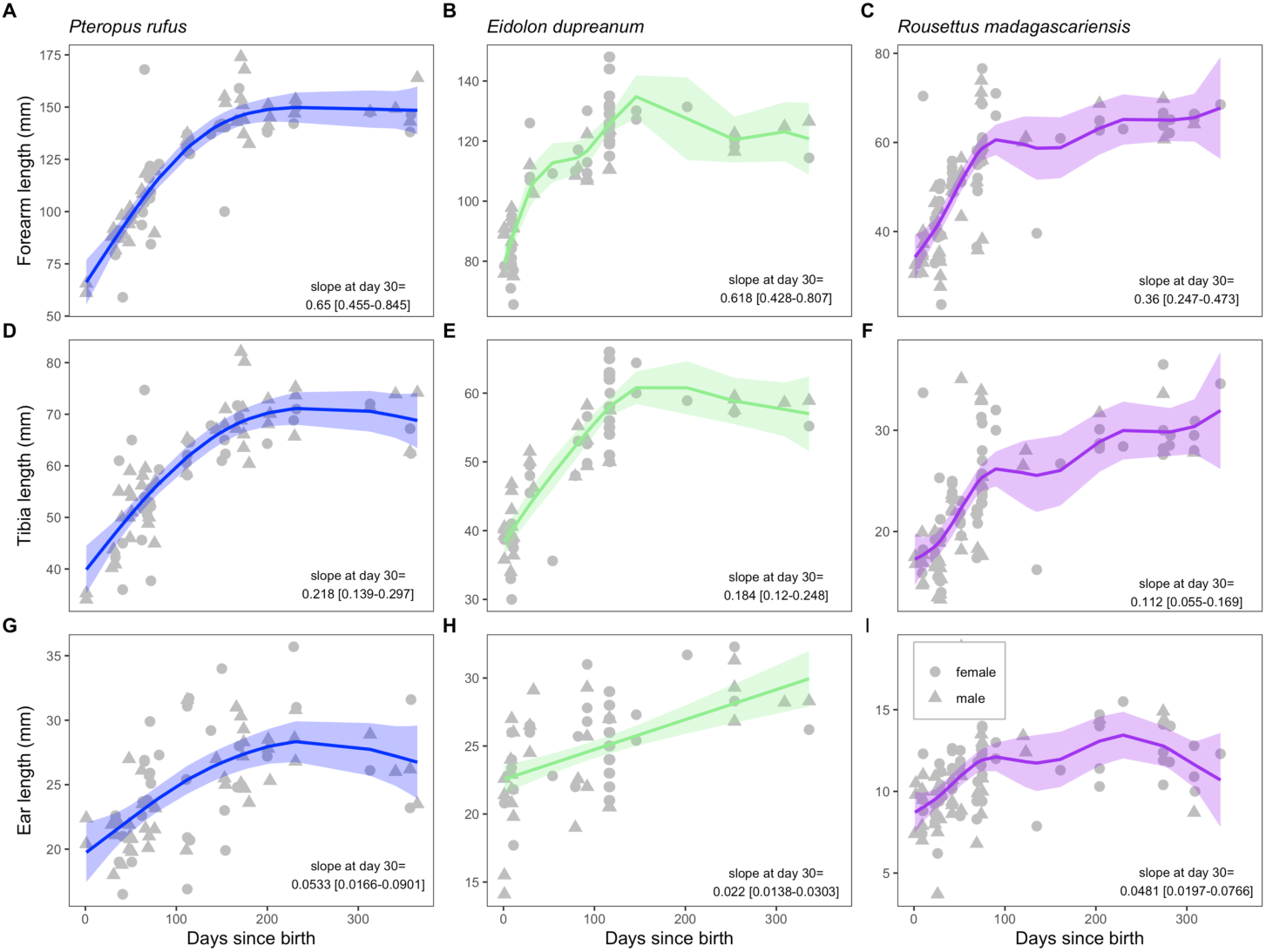
Variation in juvenile forearm, tibia, and ear length with days since birth, corresponding to the date of first observed juvenile for each of three Madagascar species (Sep-29 for *P. rufus*, Nov-16 for *E. dupreanum*, Dec-12 for *R. madagascariensis*). Raw data are shown in grey (females = triangles, males= circles), with predictions fitted GAM model overlain as colored lines; 95% confidence intervals by standard error are shown by translucent shading (Table S6). The slope (derivative) of each fitted GAM at day 30 is identified in the bottom right; derivative results across the first year are summarized in Table S7.

## Discussion

Here, we explore spatial-temporal and seasonal variation in morphological features for three endemic Malagasy bats in the Old World Fruit Bat family, Pteropodidae: *P. rufus, E. dupreanum*, and *R. madagascariensis*. Our work confirms that *P. rufus, E. dupreanum*, and *R. madagascariensis* birth in a single annual, species-specific pulse in Madagascar, which is temporally staggered across the three species. In the District of Moramanga in Madagascar’s center-east where we conducted the bulk of our field studies, the *P. rufus* birth pulse occurred first in the months of September/October, followed by *E. dupreanum* in November, and *R. madagascariensis* in December. It is possible that the timing of this birth pulse may vary latitudinally based on climatic differences across the island (e.g. occurring earlier in warmer climates or later in cooler regions), though our birth pulse projections align well with previous records of the mating season for *P. rufus* in southeastern Madagascar (Long and Racey 2007) and *R. madagascariensis* in northwestern Madagascar (Noroalintseheno Lalarivoniaina et al. 2019); to our knowledge, no previous records defining the reproductive calendar for *E. dupreanum* have been published (Shi et al. 2014). Nonetheless, climate-related variation in birth pulse timing is well-described for populations of *Eidolon helvum*, which range across the entirety of the African continent (Peel et al. 2013, 2017).

This birth timing of Malagasy fruit bats likely increases their vulnerability to seasonally-varying population pressures. In particular, fruit bats are legally hunted during the Malagasy winter (1 May – 1 September), which overlaps the gestation period observed here for all three species, but most significantly for *P. rufus*, a species already known to be experiencing severe population declines due to anthropogenic threats (Golden et al. 2014; Brook et al. 2019a). Critically, the Malagasy fruit bat lactation periods are varied in duration such that, despite staggered birth pulses, juvenile weaning occurs largely coincidentally at the onset of the peak fruiting season in the hot-wet Malagasy summer, a pattern recapitulated across numerous species of frugivorous lemur, as well (Wright et al. 2005). As a result, Malagasy fruit bat population viability will likely be sensitive to future shifts in fruiting phenology, which are predicted to accompany changing climates (Dunham et al. 2018). Importantly, our study quantifies life history traits needed to assess population viability for these species into the future (Dobson and Lyles 1989; Brook et al. 2019a).

In Madagascar, seasonally-staggered birth pulses across the three fruit bat species could support the persistence of multi-species pathogens, such as bat-borne coronaviruses, which frequently transmit and recombine amongst different species of bats that co-roost in the same caves (Hu et al. 2017). Among Malagasy pteropodids, *E. dupreanum* and *R. madagascariensis* are known to share cave roosts, sometimes with insectivorous bats, while *P. rufus* inhabits single-species arboreal roosts (MacKinnon et al. 2003). Previous work suggests that sympatric cave-roosting likely plays a role in pathogen-sharing of diverse paramyxoviruses among Malagasy bats (Mélade et al. 2016), but considerable evidence also supports a largely single-host-species-to-single-pathogen relationship for many other bat-borne infections, including those described in Madagascar (Ng et al. 2015; Lagadec et al. 2016; Brook et al. 2019b; Joffrin et al. 2019). It is likely that diverse inter- and intra-species dynamics underpin the population-level persistence of different pathogen types.

Because the dynamics of pathogen shedding and zoonotic spillover have been linked to reproductive and nutritional calendars across several bat-virus systems (Plowright et al. 2008; Amman et al. 2012; Schmidt et al. 2017; Brook et al. 2019b), documentation of seasonal variation in bat body condition and nutrition also has important implications for understanding immunity and pathogen maintenance. We here highlight significant seasonal changes observed in body condition for Malagasy fruit bats, apparently modulated by reproduction for females and corresponding more closely to the nutritional calendar for males. Further research confirming the reproductive status of adult female bats in this system—by either ultrasound in the field or assay of plasma progesterone from field-collected samples (Buchanan and Younglai 1986)—is needed to confirm this hypothesis of reproductive regulation of seasonal female bat masses. Additionally, future work elucidating seasonal and cross-species variation in fruit bat diet—and its impact on bat health—would do much to elucidate the observed discrepancy in the timing of the seasonal mass: forearm deficit for in *E. dupreanum* (June-July) vs. *P. rufus* (September) males. No seasonal pattern was found for male *R. madagascariensis* bats in our dataset, which could result from a lack of statistical power to identify differences across a smaller body size range for this species, or which may signify perpetually abundant food resources for this species in the Moramanga District.

Beyond the observed seasonality in body mass:forearm residual, which tracked reproduction for females and nutrition for males, we also documented sexual dimorphism (larger males vs. females) in tibia and forearm lengths for Malagasy fruit bats, a pattern that is common to pteropodids more generally and has been previously reported for *R. madagascariensis* (McNab and Armstrong 2001; Goodman et al. 2017). Our study confirms that the size distribution of Malagasy pteropodids spans the range of that documented globally, with *P. rufus* and *E. dupreanum* falling among the larger 50% of previously described species and *R. madagascariensis* among the smaller. Mirroring adult size distributions, juvenile growth rates were highest and developmental periods longest in *P. rufus*, followed by *E. dupreanum* and *R. madagascariensis*. Critically, the longer development period for *P. rufus* further jeopardizes this species’ already-threatened conservation status—which recent analysis suggests may be even more vulnerable than previously reported (Brook et al. 2019a). The rapid ∼two month juvenile growth window witnessed for *E. dupreanum* in our dataset suggests that this species may actually birth earlier than is recorded here; additional, intensive sampling throughout the reproduction period is needed to confirm the seasonal limits of each developmental stage for these three fruit bat species.

In conclusion, we quantify life history traits needed for population modeling, document seasonal variation in body condition, and elucidate the Malagasy fruit bat reproductive calendar, contributing important resources for future efforts to quantify both conservation trajectories and zoonotic pathogen transmission in his system. This work emphasizes the importance of longitudinal field studies in uncovering seasonal variability in ecological data, with critical implications for understanding of both population viability and infectious disease dynamics alike.

## Acknowledgements

The authors thank Kimberly Rivera, Katie Fitzgerald, and Samantha Kreling for help in the field, the Virology Unit at the Institut Pasteur de Madagascar for logistical support, the Department of Zoology and Animal Biodiversity at the University of Antananarivo for help in obtaining research permits, and the Brook lab at the University of Chicago for helpful contributions to the manuscript. We acknowledge funding from the National Institutes of Health (1R01AI129822-01 grant JMH and CEB), DARPA (PREEMPT Program Cooperative Agreement no. D18AC00031 to CEB), the Adolph C. and Mary Sprague Miller Institute for Basic Research in Science (postdoctoral fellowship to CEB), and the Branco Weiss Society in Science (fellowship to CEB).

## Appendix 1: Supplementary Figures

**Fig. S1.**
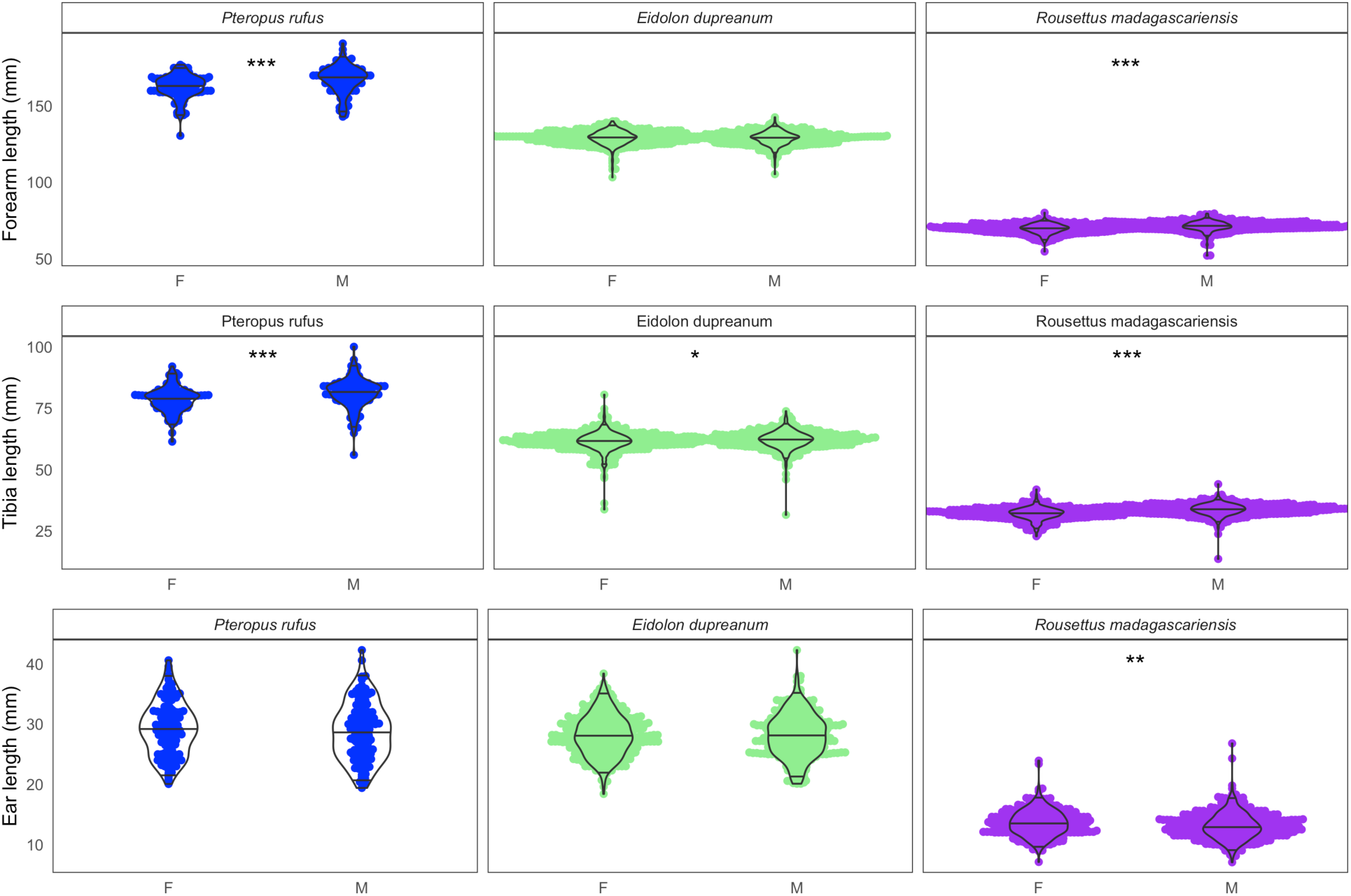
Morphological trait distributions in forearm, tibia, and ear length for *Pteropus rufus, Eidolon dupreanum*, and *Rousettus madagascariensis*. Each panel compares distributions from females (F) vs. males (M). Asterisks indicate significance in *Welch’s 2-sample t-tests* from data within each panel, based on significance codes: ***=0.001; **=0.01; *=0.05;. = 0.1.

**Fig. S2.**
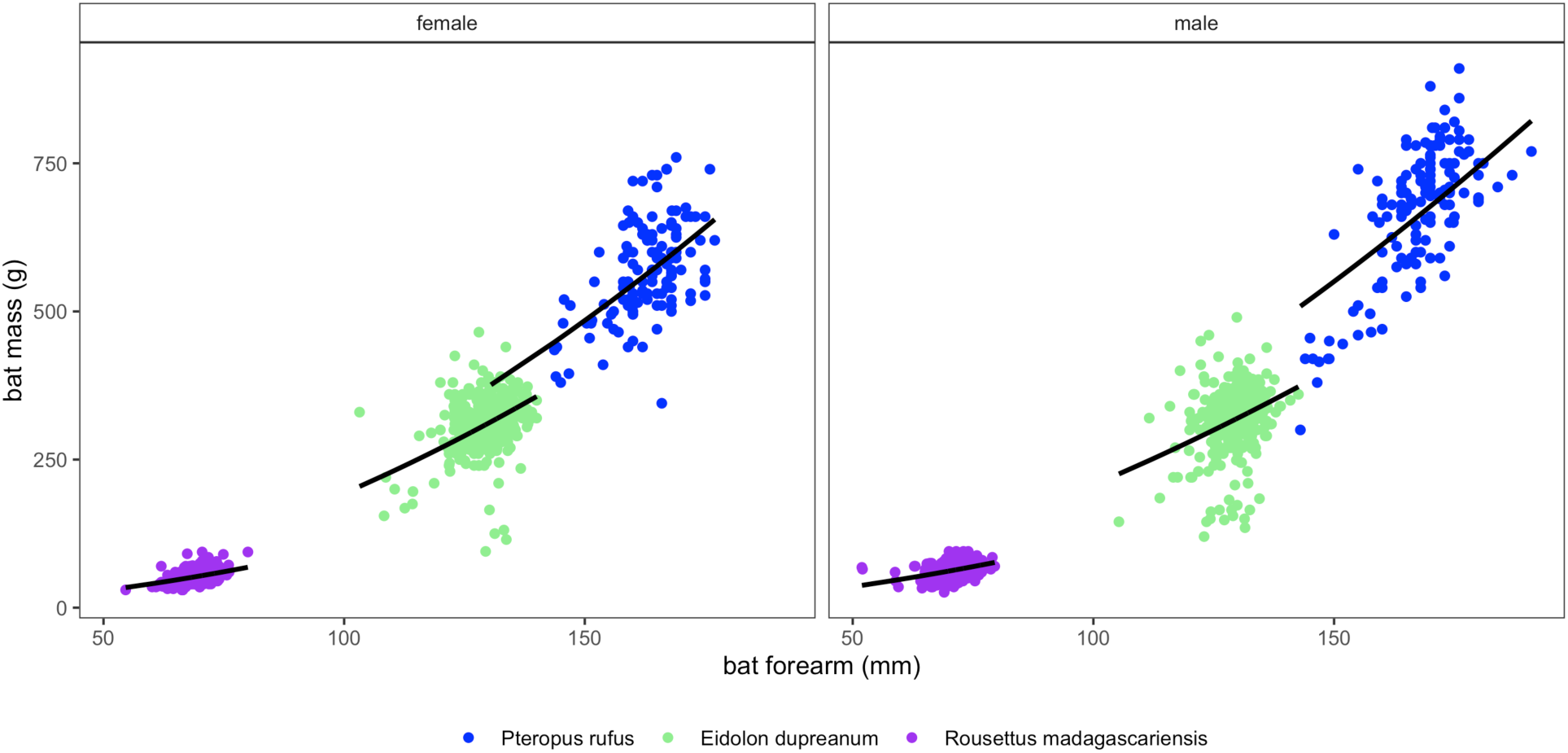
Forearm length (mm) vs. mass (g) relationships for Malagasy fruit bats, separated by sex. Data are depicted as points colored by species. Solid black lines correspond to output from fitted linear regression model of log10 mass predicted by log10 forearm length, incorporating a fixed predictor of species. Residuals depicted in Fig. 3 (main text) were derived by subtracting predictions from data for each individual, as shown here. Model fits are summarized in Table S4.

**Fig. S3.**
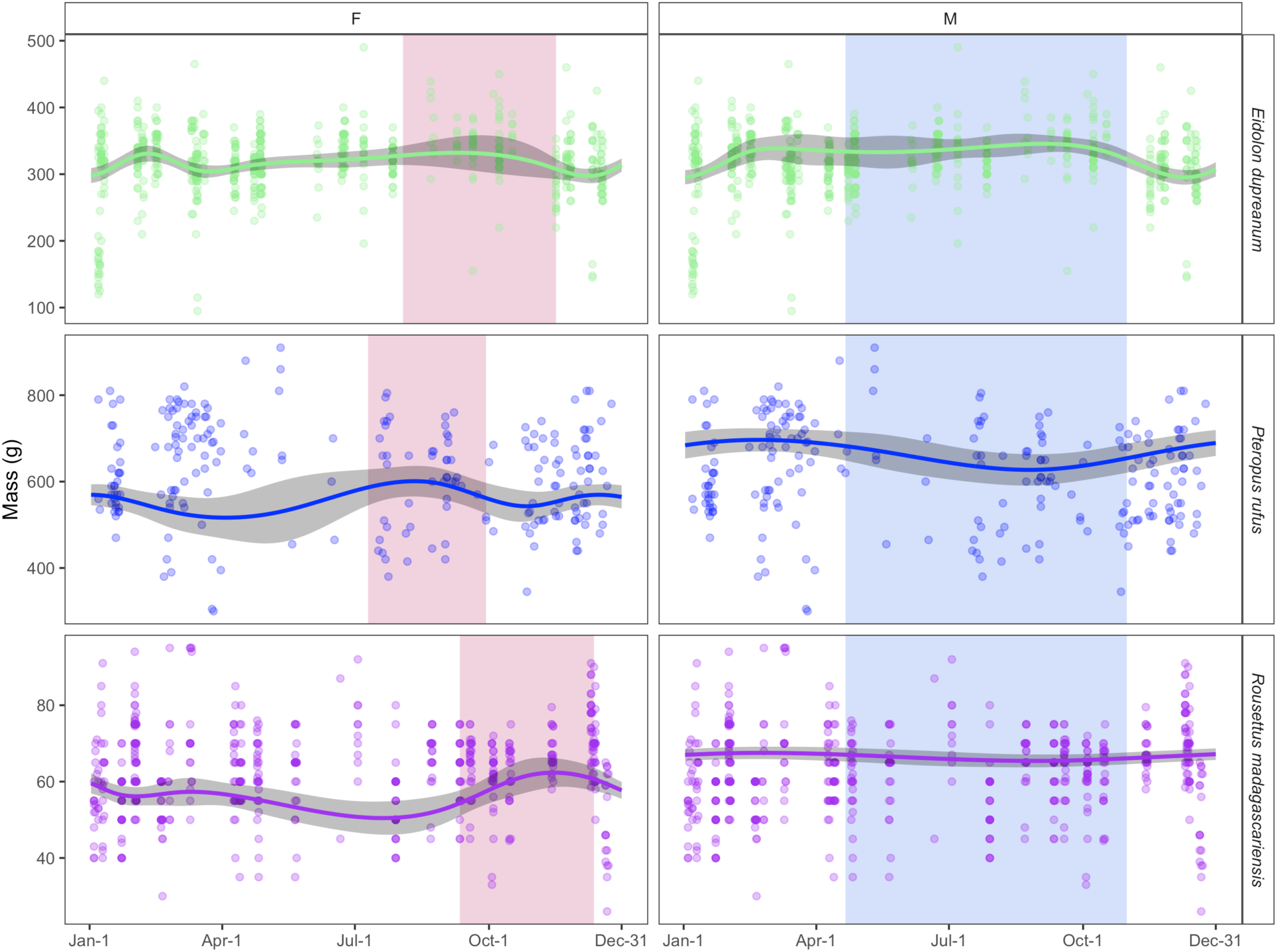
Figure largely replicates Fig. 3 (main text) but here depicts the output of supplementary GAMs incorporating a direct response variable of bat mass (in g) predicted by day of year (as a cyclic cubic smoothing spline) with a random effect of forearm length. Random effects are silenced here for plotting purposes. As in Fig. 3, raw data are shown as open circles with prediction from fitted GAM model as solid line; 95% confidence intervals by standard error are shown by shading in gray (Table S4). For female plots, pink shading corresponds to the species-specific gestation period; for male plots, blue shading corresponds to the winter dry season in Madagascar.

